# Autogenic spinal excitatory circuit ensures skilled hand movements in primates

**DOI:** 10.1101/2025.03.31.646263

**Authors:** Jihee Kim, Saeka Tomatsu, Tatsuya Umeda, Tomohiko Takei, Tetsuro Funato, Kazuhiko Seki

**Affiliations:** Department of Neurophysiology, National Institute of Neuroscience, Tokyo 187-8502, Japan; Department of Developmental Physiology, National Institute for Physiological Sciences, Aichi 444-8585, Japan; Department of Mechanical Engineering and Intelligent Systems, The University of Electro-communications, Tokyo, Japan

## Abstract

Skillful hand movements are a hallmark of primates, including humans, requiring sophisticated motor planning and execution. Challenging the longstanding paradigm that emphasizes the dominance of the cerebral cortex^1^, our study highlights a pivotal role for spinal excitatory reflex circuits in both planning and executing skillful hand movements. Using a combination of experimental approaches with behaving non-human primates and predictive simulation, we identified a group of excitatory spinal interneurons that orchestrate a closed-loop, positive feedback mechanism during voluntary wrist movements. This mechanism is characterized by a bidirectional interaction between inter-neuronal spiking and muscle activity, mediated by motoneuronal efferent signals and proprioceptive afferent signals from the same agonistic muscles. Furthermore, we demonstrate that the temporal profile of muscle activity during movement execution, including amplitude and duration, is pre-determined during motor planning at the spinal interneurons, functioning as a force-feedback gain within the excitatory circuit. These findings suggest that autogenic spinal excitatory circuits play a predominant role in shaping overall muscle activation during motor execution, provided the proper reflex gain is pre-set by higher neural systems during motor planning. Together, our findings provide a cellular-level perspective on how spinal reflex loops contribute to skilled voluntary movements in primates, extending over a century of spinal reflex research^2^.

## INTRODUCTION

The spinal cord, which is intricately connected to the brain, plays a crucial role in coordinating various body movements, and is especially significant in controlling stereotyped actions, such as locomotion and postural control. Within the spinal cord, segmental reflex circuits constitute the core modules for sensorimotor transformations during movement. These reflex circuits, present in diverse species, from invertebrates to higher mammals, including humans, directly, efficiently, and quickly integrates somatosensory signals into ongoing motor actions. Physiological studies^3,4^ and recent advancements in genetic tools^5,6^, have led to the characterization of several reflex circuits and their key interneurons, delineating their distinct input–output properties. Despite extensive evaluation in relation to stereotyped movements^7^, the involvement of these circuits in the execution of non-stereotypical movements^8^, such as skilled voluntary movements, remains largely unexplored. Traditionally, the cerebral cortex and corticospinal tracts have been widely considered the primary drivers of muscle activation for executing primate hand actions, such as grasping^1,9–12^. By demonstrating the minimal effects of motor cortex lesions on the execution of learned and skilled motor actions in non-primate models, in cat^13^, rat^14–16^, and mouse^17^, several studies have challenged previous theories of exclusive cortical function. Despite anatomical differences between primates and other species that may underlie this discrepancy^18^, a fundamental question arises of the mechanism mediating muscle-activity control with the limited involvement of the cerebral cortex in primates, including humans.

Here, we posited that the spinal reflex circuit is potentially an alternative driver of muscle activation during voluntary movement in non-human primates. This hypothesis was investigated based on the spinal excitatory interneuronal activity that mediates voluntary muscle activation in behaving monkeys, and we modelled their implementation in the neural system that controls voluntary motor action.

## RESULTS

We trained three monkeys to perform wrist flexion-extension tasks ^19^ (Extended Data Fig.1), which constitute non-stereotyped sensorimotor behavioural tasks because the monkeys need to adjust the kinematics and dynamics of their wrist movements according to varying visual instructions. During the performance of the trained task, we recorded the activity of interneurons (IN) which mediate the segmental spinal reflex (Reflex IN) in the cervical spinal cord. The reflex INs were identified using both input from and output to the periphery. Based on the response to the electrical stimulation of the primary sensory (Group I) afferents of the wrist extensor muscles (deep radial nerve (DR), Fig. 1A, B), input was examined^19,20^ whereas the output was determined by spike-triggered averaging (STA) of reflex IN’s effect on the motoneuron pool of the wrist and finger muscles (Fig. 1C, D)^21,22^. Of the 292 neurons recorded in the monkeys, 35 reflex INs were identified (Extended Data Fig. 2A) and, based the target muscle of the reflex neurons, were categorized into six possible input–output patterns (Extended Data Fig. 2B). These reflex INs predominantly evinced an ‘autogenic’ pattern (i.e., INs receiving Group I afferents from wrist-extensor muscles and also projecting to the wrist-extensor motoneurons, n=20/35), and a majority (75%) were excitatory IN (Extended Data Fig. 2B). Excitatory INs that mediated autogenic reflexes (n=15) were named ‘agINs’ (Fig. 1E). The representative activity of agINs during voluntary wrist movements (Fig. 1F, Extended Data Fig. 1) is shown in Fig. 1G (wrist extension) and 1H (flexion; the same neurons shown in Fig. 1B and 1D). The agIN firing pattern was highly biased towards wrist extension (Fig. 1I). Among the 15 agINs, 13 showed extension-biased activity, one showed flexion-biased activity, and one showed an unbiased activity pattern, indicating a highly significant bias toward extension (*p*<0.001, binomial test); the ground averages of these 15 agINs also exhibited extension-biased activity (p=0.002, two-way ANOVA, Fig. 1J).

**Figure 1.**
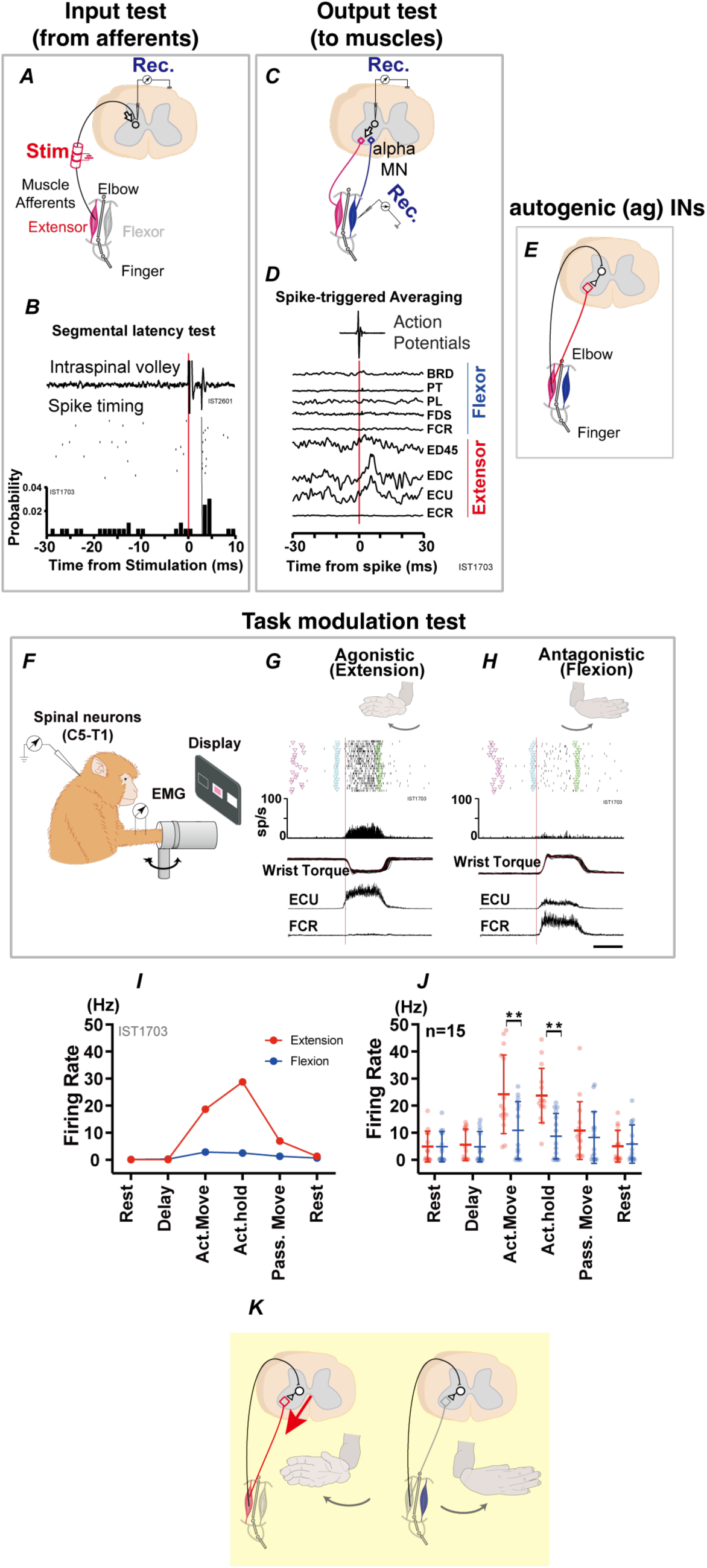
Excitatory interneurons mediating autogenic facilitation of wrist-extensor muscle. **A-E**: Identification of interneurons that mediate segmental spinal reflexes. **A and B**: Examination of afferent input and **Cand D** of output projection to the wrist motor neuron. **A**: The afferent nerve innervating the wrist extensor muscles (deep radial nerve; DR) was stimulated using an implanted nerve cuff electrode (Stim), and neuronal responses as well as an incoming volley were recorded (Rec) in the C6-T1 segments in monkeys performing the task. **B**: A representative neuron showing a monosynaptic response to stimulation. Neurons showing response within a central latency of less than 1.5 ms (from the volley) were deemed as the first-order interneurons. **C**: For the neurons showing monosynaptic response to DR stimuli, their output connection to motoneurons of the wrist flexor and extensor muscles was determined by averaging electromyographic (EMG) signals triggered by the timing of each action potential of the interneuron. **D**: pre- and post-spike effects of the same neuron as B. From the input (B) and output (D) profile, this example neuron was identified as the excitatory interneurons mediating afferent input (from DR afferent) and projecting to the motoneuron (EDC and ECU) of both wrist extensor muscles (“agINs,” E). BRD, brachioradialis; PT, pronator teres; FDS, flexor digitorum superficialis; FCR, flexor carpi radialis; ED45, extensor digitorum-4,5; EDC, extensor digitorum communis; ECU, extensor carpi ulnaris; ECR, extensor carpi radialis. **F-H**: Evaluation of the firing pattern of the identified interneurons in a monkey performing voluntary movements. While the monkeys performed the wrist flexion and extension tasks with an instructed delay period (Extended Data Fig. 1), the activity of the interneurons was recorded (F). **G, H**: Example of the activity of reflex interneurons (the same neurons as B and D) during extension (agonistic to the DR nerve, G) and flexion (antagonistic to the DR nerve, H) trials. From top to bottom: dot-raster plot, peri-event time histogram, torque at the wrist (+: flexion, -: extension), ECU EMG, and FCR. Each sweep was aligned with the onset of wrist torque (red line). Data from 21 flexion and extension trials are presented. **I-J**: Modulation of the firing rates of reflex INs during the task. I: Mean firing rate in each task epoch of the same interneurons B, D, G, and H. Delay: Instructed delay period, Act. Move: Active movement; act. Hold: Active hold period; Pass. Move: passive movement period. **J**: Mean±SD of the firing rate of all agINs (n=15; Extended Data Fig2). **; *p*<0.01. K: Schematic description of the input-output relationship of agINs and corresponding firing pattern during wrist extension (left) and flexion (right) tasks.

Based on the characteristic input-output pattern of agINs, they could be identified as the ‘Ib INs’^4,23^, which mediate the Group I oligosynaptic-^24^ or disynaptic^25,26^ input to the extensor motoneuron from both Ia and Ib afferents^4,27,28^. In ‘reflex reversal’, the Ib INs became excitatory during locomotion^25,29^, whilst otherwise suppressing synergistic motoneurons^30^. We hypothesized that the agIN’s predominant activity during extension reflects their unique role in controlling voluntary movements by generating and modulating agonistic muscle activity (Fig. 1K) through Group I afferents^31^, which are activated via a recurrent positive-feedback reflex loop involving the same agonistic muscles.

We tested this hypothesis by examining whether (1) the activity of the extensor muscle during wrist extension effectively influences the activity of agINs (Fig. 2A, *decoding*), and (2) the activity of agINs can generate extensor muscle activity (Fig. 3A, *reconstruction*).

**Figure 2.**
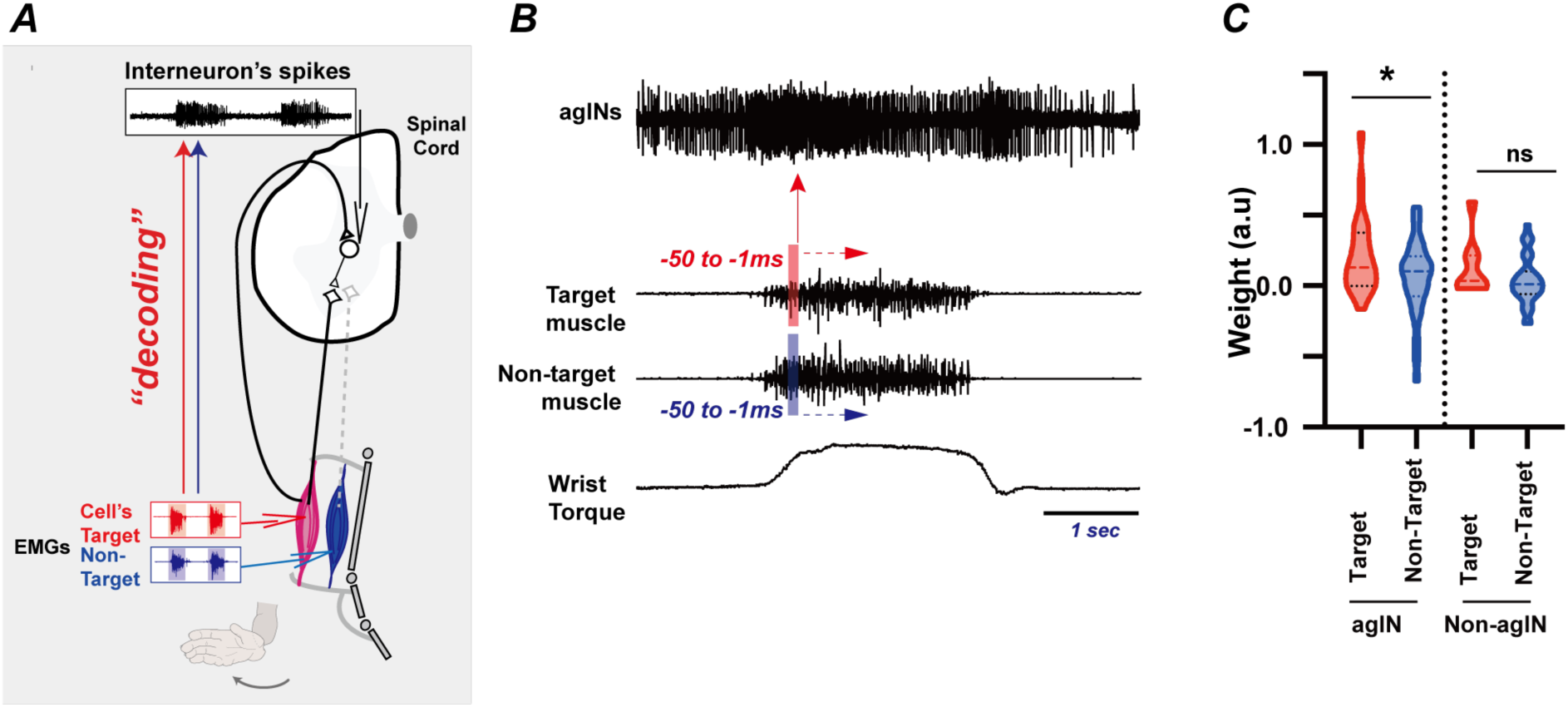
Decoding the agIN firing pattern based on the activity of their target muscles. **A**: The supposed closed loop consists of agINs, its target extensor motoneuron (black diamonds) and muscle (red), and sensory afferent fibers directly projected to the same agINs. Non-target agonistic motoneurons (grey diamonds) and muscles (blue) are also illustrated for comparison. The capability of the activity of agINs’s target muscle to affect the spiking activity of agINs during agonistic movement (bottom, wrist extension) was examined by comparing the performance of ‘decoding’ the agINs activity by the EMG of the target muscle (red) and non-target muscle (blue). **B**: Details of the decoding analysis. For each agIN, the influence of the extensor muscle activity on its future spiking activities (≤50 ms) was examined by comparing the decoding performance using the EMG of the target muscle (red arrow) and that of the non-target muscle (blue arrow). See the details of the methods. **C**: Left, weight value of the EMG of the target (red) and non-target (blue) extensor muscles in models that predicted agIN activity. *; *p*<0.05. (Unpaired *t*-test). Right: same comparison in non-agINs.

**Figure 3.**
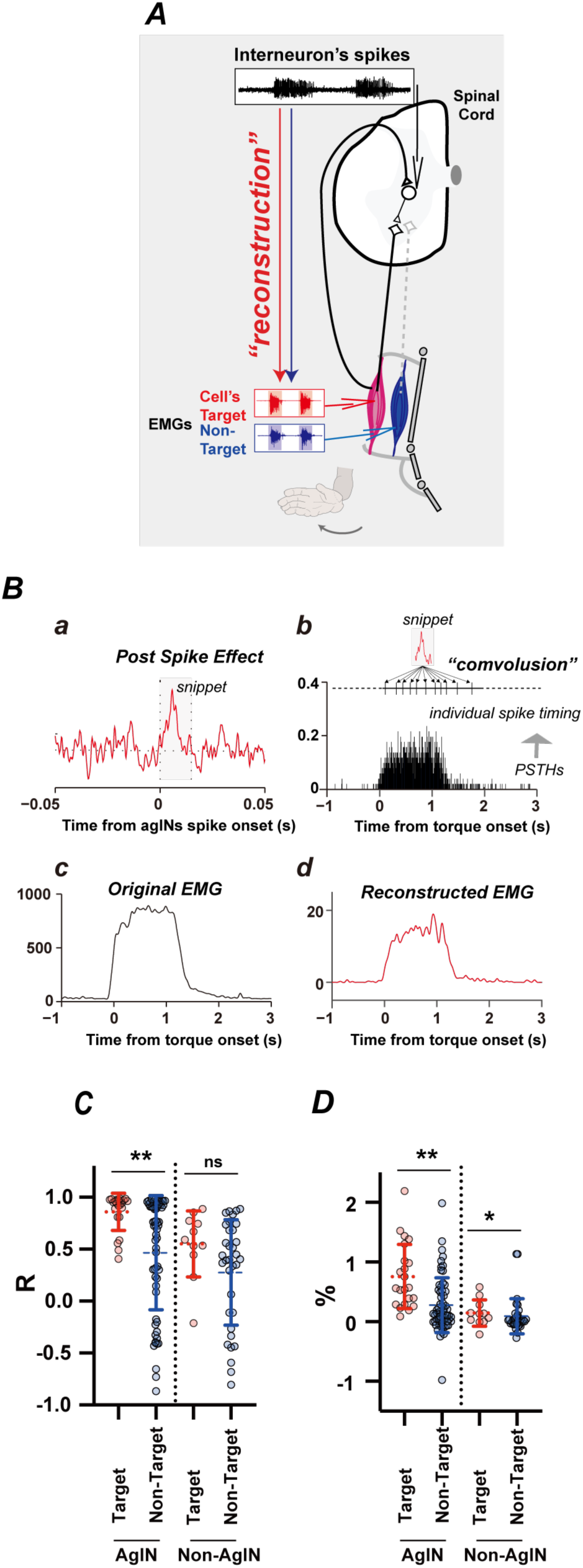
Reconstruction of the muscle activity by the post-spike effects and the firing pattern of the agINs. **A**: The contribution of the agINs to generating the EMG of their target muscle was examined by the ‘reconstruction’ analysis. **B**. Reconstruction analysis to evaluate the contribution of single agINs in generating the EMG signals of target muscles. ***a***: Waveform of post-spike facilitation of the target muscle (ECU) for single agINs. Shading represents the snippet applied for this analysis (15ms from the trigger). ***b***: The firing pattern of the same agINs as ***a*** is aligned by the movement onset). Bin width=5ms. Compiled from 51 extensor trials. **c**: EMG of the target muscle (same muscle as **a**). Smoothed and averaged for 51 extensor trials. **d**: Reconstructed EMG generated by **a** and **b**. Note the difference in the vertical scale between ***c*** and ***d*** *(see main text)*. **C**. Left, Correlation coefficient between the original EMG (as in Bc) and reconstructed EMG (as in Bd) of the target (red) and non-target (blue) muscles of the ag-INs. Means(dotted lines) ±SD (continuous lines). **; *p*<0.01. Right: same comparison in non-agINs. **D.** Comparison of the size of the reconstructed EMG relative to the original EMG shown in the same format as in panel C.

First, using electromyographic (EMG) signals of the wrist muscles, we decoded the agIN firing activity during wrist extension. This analysis focused on the 50- to 1-ms duration preceding the occurrence of each agIN action potential (Fig. 2B) with the rationale that, if agIN firing is partly driven by sensory afferent signals from their target muscles, a higher decoding weight could be expected using EMGs of their target muscles (red arrow in Fig. 2A, left), rather than of the synergistic, non-target muscle (blue arrow). As each agIN receives a direct projection from the extensor afferent (Fig. 1B), this analysis could determine whether the extensor muscle’s activity drive agINs through this projection. The agIN firing profile was successfully predicted by the EMG of the target muscle, both at the single-test base (Extended Data Fig. 3A) and population level (Extended Data Fig. 3B). Furthermore, using the agIN’s target muscle (n=22) instead of that of their non-target synergistic muscles (Fig. 2C left, n=29, *p*=0.045, unpaired *t*-test), we found a higher decoding weight in the pair. A comparable result in the non-agINs (i.e., reflex IN with post-spike effect on the flexor muscles, ‘heterogenetic’ and ‘both’ neurons in Extended Data Fig. 2) between the target and non-target muscles (Fig. 2C right) was not identified, which suggests that the observation is agIN specific. As the transcortical sensorimotor loop cannot influence agIN activity within the time range for decoding (∼50 ms^32^), we infer that the agIN is directly driven by reafference from its target muscles.

Next, we examined the contribution of agINs to generating the activity of their target extensor muscles during wrist extension (Fig. 3A, the arrow ‘reconstruction’) ^33^. First, we compiled a snippet (Fig. 3Ba) from the STA of both target and non-target muscles that reflects the contribution of a single spike of agINs to generate muscle activity^34,35^. By convolving each snippet into the timestamp of the agIN spikes (Fig. 3B*b*), we reconstructed the EMG signals of the target muscle, averaged them over each trial (Fig. 3B*d*), and compared them with the original EMG signal of the same muscle (Fig. 3B*c*). In this example (See Extended Data Fig. 4 for another example), the reconstructed EMG exhibits a temporal profile that correlates with the original EMG (from onset to offset; n=10,383 points, R=0.966), and further compared the magnitudes (total area from onset to offset of the original EMG) of the reconstructed and original EMG signals by calculating their ratios. The ratio in the example (1.55% of the total area of the original EMG) was greater compared to those of three non-target extensor muscles (n=3) of this agIN (0.39±0.28% (SE) [0.10, −0.005, 1.09]).

We repeated the same analysis for all 22 pairs of agIN target muscles. The similarity with the original EMG profile was higher when the reconstructed EMG was created from the snippet of the target muscle (R=0.86±0.03, n=22) than of non-target muscles (R=0.46±0.07, n=61, *p*<0.01, unpaired *t*-test, Fig. 3C, left). We found the non-agIN showed much lower similarity of the reconstructed EMG with their target muscles [Fig. 3C, red on the right and left, R=0.86±0.03 (agINs) vs. 0.55±0.09 (non-agINs), n=22 vs. n=11, *p*<0.01, unpaired *t*-test]; furthermore, the non-agINs showed indifferent similarity in the EMG reconstructed either by their target or non-target muscles [Fig. 3C right, R=0.55±0.31 (target muscles) vs. 0.27±0.50 (non-target muscles), n=11 vs. n=34, *p*=0.10, unpaired *t*-test]. We also found that the single agIN could generate 0.75% (±0.12%) of the original EMG, which is larger than the reconstructed EMG of non-target muscles (Fig. 3D left, 0.27±0.05%, n=22 vs. n=61, *p*<0.001, unpaired *t*-test). Thus, if we assume that only agINs generated the EMGs, we infer that the original EMGs of the agIN target muscle could be generated by only 114 to 158 (on average, 133) agINs. This number represents the exclusive capability of agINs for motoneuronal activation, because thousands of spinal^36^ and non-spinal^37,38^ premotor neurons could mediate EMG generation ^33^. In contrast, the reconstructed EMGs of the non-agINs’ target muscles were smaller [size=0.75%±0.11% (agINs) vs. 0.14%±0.06%, n=22 vs. n=11, *p*<0.01, unpaired *t*-test), and showed little differences between the non-agINs’ target or non-target muscles (Fig. 4D right, 0.14%±0.06% vs. 0.09%±0.05%, n=11 vs. n=34, *p*=0.58, unpaired *t*-test). These results suggest that agIN mediates online muscle-activity augmentation during voluntary movement through a positive-feedback reflex loop.

**Figure 4.**
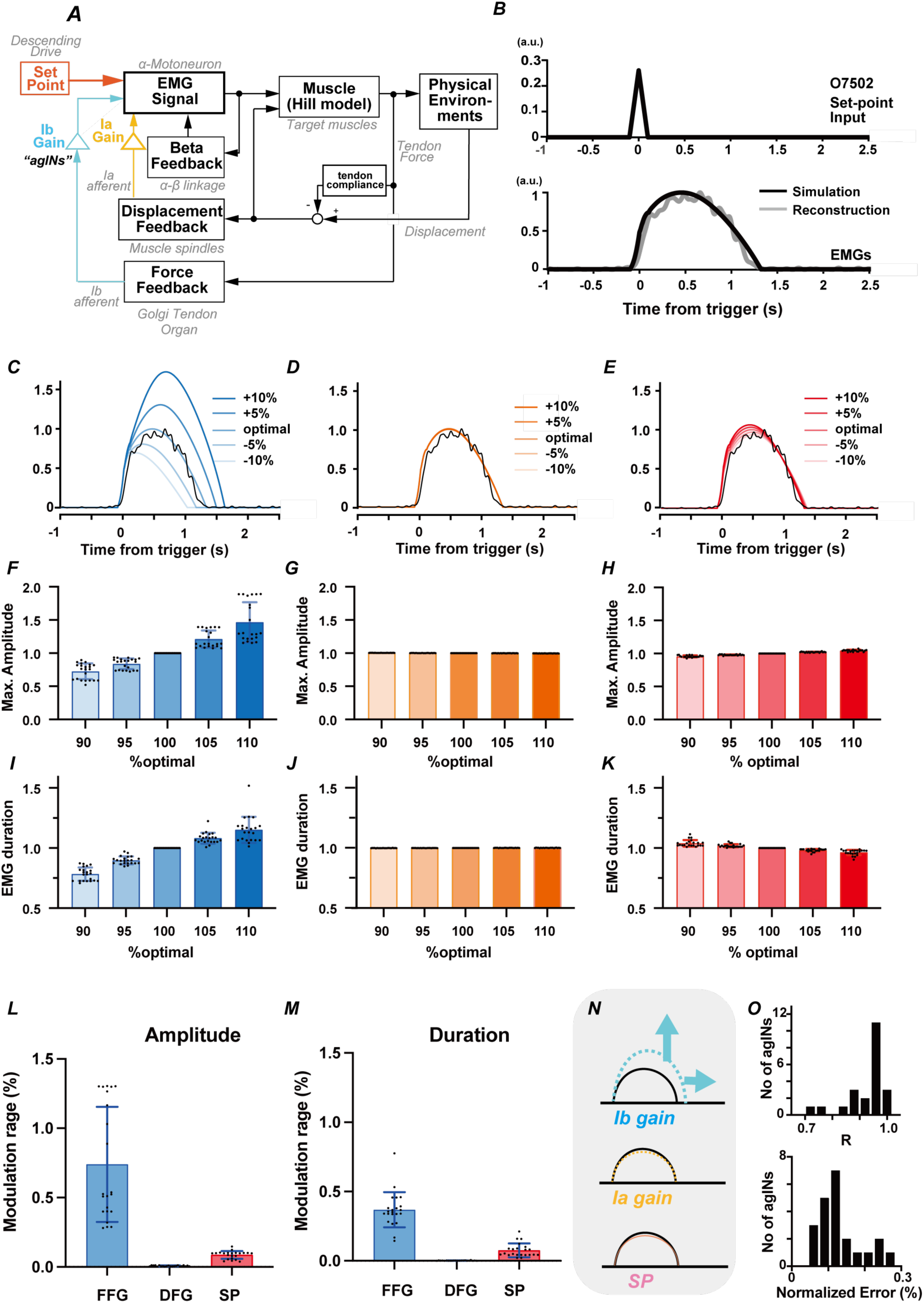
Simulation. **A**: Simplified block diagram of the nonlinear model of segmental reflex system mediated by the agINs which is modified from the existing segmental reflex system model ^39^. All model parameters, other than the gain parameter, were fixed to the value in the previous report ^78–81^. See methods for details. The activity of agINs recruit and augments alpha motoneuron activity and generates EMG in the target muscle. Through the contractile properties and physical environment, the EMG generates autogenic force feedback (via Golgi tendon organ and Ib afferents) and reciprocal displacement feedback (via muscle spindle and Ia afferent) that recurrently modulate the activity of alpha motoneurons and target muscles. This model assumes the agIN is driven by the autogenic force feedback ^25,29,78^. **B**: A reconstructed (grey, n= 89 trials) and simulated (black) EMG of wrist-extensor muscle (ED23) in the extension trial. Simulated EMG was obtained by using the optimal FFG (1.742), DFG (0.871), SP (0.260). R= 0.99. **C-E:** Example of the effect of different FFG (blue, **C**), DFG (orange, **D**), or SP (red, **E**) on the simulated EMG waveform. **F-H:** the effect on the maximal amplitude of simulated EMG waveform (n=21). Note that one neuron-muscle pair has been eliminated from this analysis because of the inability to define the sustaining duration due to the lack of clear offset in the reconstructed EMGs. **I-K**: the effect on the duration of simulated EMG waveform. **L-M**: effects of changing the FFG, DFG, and SP from 90 to 110% of their optimal value on the amplitude (**L**) or the duration (**M**) of reconstructed EMGs. Mean (bar) and SD (line) were also plotted. **N:** A proposed regulation of EMG burst duration and amplitude by the agINs’ loop. The FFG affects both the amplitude and duration of upcoming EMG burst; the DFG nor SP had little effect. **O**: Distribution of the correlation coefficient (top) and normalized difference (bottom) between the simulated and reconstructed EMGs. n=21.

Next, we simulated how the agIN-mediated positive-feedback reflex loop was implemented during muscle activation. In an existing model of the segmental reflex system^39^, we simulated the EMGs reconstructed from each agIN (Fig. 4a). Based on the assumption (see Fig.1) for the identity of agINs as the ‘Ib interneuron’^4,23^, which predominantly receives direct input from tendon receptors, we assume that agINs are solely activated by force-feedback signals from the Golgi tendon organ of the agonist muscle. A remarkable feature of this model is that it can generate time-varying EMG signals that simulate actual reconstructed EMGs with higher accuracy (R=0.99, Fig. 4b) by setting several fixed numerical parameters.

First, we systematically changed the force-feedback gain (FFG; Ib gain, blue in Fig. 4A), displacement-feedback gain (DFG; Ia gain, orange in Fig. 4A), and the amplitude of the setpoint input (SP; red in Fig. 4A), whereas the other model parameters were fixed (Extended Data Table 1). Representative results are shown in Fig. 4C–E (data from the same agINs as in Fig. 4B are used). When we increased and decreased the FFG (Fig.4C) around the optimal FFG (providing the best match between the simulated and reconstructed EMG signals), we observed larger and smaller EMG signals (amplitude, n=5; R=0.94) with a longer and shorter duration (n=5; R=0.91). In contrast, altering the DFG (D) or SP (E) conferred minimal impact on the simulated EMG signals for amplitude (DFG; n=5, R=1.0, SP; n=5, R=1.0) or duration (DFG; n=5, R=0.87, SP; n=5, R=0.96). The same procedure was repeated for 21 agIN–muscle pairs. The results are summarized in Fig. 4F–K. When the FFG was increased from 90% to 110% of the optimal value, both the magnitude (Fig. 4F) and the duration (Fig. 4I) significantly increased (G; F=15.94, *p*=0.0006; 0.68±0.17 (90% optimal) to 2.22±1.59 (110% optimal), *p*=0.0015 with Dunnet’s multiple-comparison test, J; F=10.56, *p*<0.002; 0.88±0.15 to 1.02±0.07, *p*<0.011). Increasing either DFG (G, J) or SP (H, K) induced subtle or little alternation (DFG, amplitude: F=16.95, *p*=0.0005; 1.00±0.00 (90% optimal) to 0.99±0.00 (90% optimal), *p*=0.0018, duration: F=1.59, *p*=0.22; 1.00±0.00 to 1.00±0.00, *p*=0.53, SP, amplitude: F=19.06, *p*=0.003; 0.96±0.04 to 1.04±0.04, *p*=0.001 with Dunnet’s multiple-comparison test, duration: F=9.76, *p*=0.0006; 1.01±0.01 to 0.99±0.01, *p*=0.0016). The effects of changing the parameter from 90% to 110% of their optimal values were larger in the FFG than in DFG or SP in both the amplitude (Fig. 4L, F=15.03, *p*=0.0009; *p*=0.0018 (with DFG) or *p*=0.0019 (with SP) with Dunnet’s multiple-comparison test) and duration [Fig. 4M, F=12.47, *p*=0.0019; *p*=0.0025; with DFG or *p*=0.0063 (with SP)] of the reconstructed EMGs. This simulation suggested the possibility that the EMG output from the agIN-mediated spinal reflex circuit could be regulated more efficiently by changing the FFG (Fig. 4N).

Second, we examined how well the simulation using the optimal FFG predicted the reconstructed EMG signals. As shown in Fig. 4B, we observed a high similarity between the actual and simulated EMGs with optimal FFGs (FFG=1.867, R=0.99). We found (Fig. 4O) higher correlation coefficients (top, R=0.91±0.08, mean±SD) with subtle prediction error (bottom, 0.13±0.06% of reconstructed EMG) between the simulated and reconstructed EMG signals from agIN (n=22). Overall, our model can sufficiently predict the EMG signals of agIN’s target muscles by tuning mostly the single-parameter FFG at the optimal value.

We repeated the same simulation to predict the actual EMG when the agIN activity was recorded (Fig. 1G and H), but not the EMG reconstructed from each agIN by changing the FFG, DFG, or SP (Extended Data Fig.5). The results were similar to those of the reconstructed EMGs, indicating that the amplitude and duration of the actual EMGs during the task can be sufficiently simulated by changing the FFG (ED Fig.4A to N) with higher precision (ED Fig.4O). Taken together, the results of these simulations suggest that the EMG output (amplitude and duration) can be regulated by modulating the force-feedback gain of the spinal reflex circuit.

Finally, we confirmed that the input–output gain of agINs (corresponding to the FFG in the simulation) is related to the size and duration of EMG in future actions during voluntary movement. We estimated the trial-by-trial changes in the FFG for each agIN by analysing the response probability of the agIN to electrical stimuli applied to the muscle afferents of the wrist extensor (Fig. 1B, see Confais et al. 2017^20^) before movement initiation (Rest, Cue, and Delay; Extended Data Fig. 1). We considered the response probability, which could be influenced by both pre- and postsynaptic factors that affect agINs^20^, as an indicator of FFG.

Our simulation (Fig. 4C, F, and I or N and Extended Data Fig. 5C, F, and I or N) indicates the possibility that trials with a high feedback gain to the ag-IN may induce a large, long EMG burst. To test this possibility, we compared the normalized amplitude and duration of the EMGs of the target muscles between trials with higher and lower afferent feedback gains. As predicted by the simulation, trials under higher afferent feedback gains exhibited larger amplitudes and longer durations of prospective EMGs (bin > 0.9, binomial test *p* = 0.003 and 0.027, Fig. 5A, B) than the baseline probability (dotted line=0.1). In contrast, no statistical bias was observed in trials with lower afferent feedback gains (bin > 0.9, binomial test p = 0.119 and 0.768, Fig. 5C, D).

**Figure 5.**
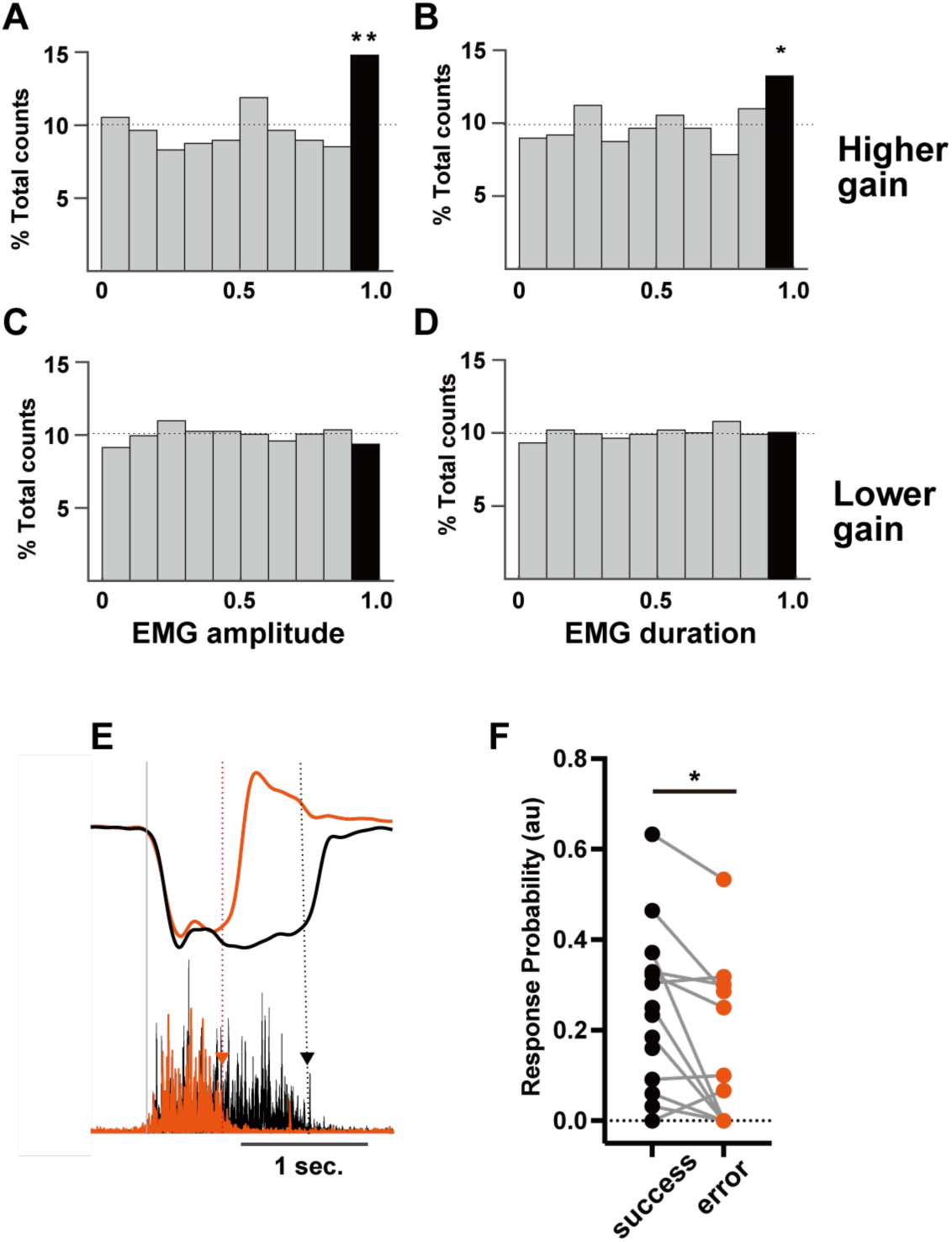
Afferent input gain to agINs and the amplitude and duration of prospective EMG burst in behaving monkeys. **A-B**: Histogram of normalized EMG amplitude (A) and duration (B) of trials with higher FFG (* *p*<0.05, ** *p*<0.101, binomial test). **C-D**: Histogram of normalized EMG amplitude (C) and duration (D) of trials with a lower FFG. **E**: Example of a single successful (black) and erroneous (orange) trial. Top: Wrist torque; Bottom: EMG of the agIN’s target muscle (EDC). Gray line: Movement onset. Black and Orange dotted lines and triangles indicate EMG offset timing. **F**: Difference in afferent input gains between successful (black) and erroneous (orange) trials. *, *p*<0.05.

Sometimes, monkeys failed the trial when they applied a shorter-duration torque (short-hold, Fig. 5E, orange) compared to the successful trial (Fig. 5E, black), even though they fully understood the rule of the task. Our simulation predicted that such trial failure might be mediated by a lower FFG. When we specifically compared the response probability of each agIN in the error trials and in the successful trials, we found the smaller FFG before unsuccessful short-hold trials in 75% of agINs (9/12; Fig. 5F) and the difference was statistically significant (Fig. 5G, 0.25±0.19 vs. 0.15±0.18, *p* = 0.0197, paired *t*-test).

Overall, these data analyses of the physiological experiments indicate that regulating the FFG of the segmental reflex pathway plays a functional role in controlling muscle activity and task performance during voluntary wrist movements, which is consistent with the prediction made by the simulation.

## DISCUSSION

The results of the experiments (Figs. 1–3 and 5) and the simulation (Fig. 4) suggest that the positive-feedback reflex system could be a predominant source for generating muscle activities during voluntary movement. Here, we assumed a simple segmental reflex circuit highlighted by agINs (Fig. 6-①), which project to the motor neurons (MN) and muscles, and the afferent muscles recurrently activate these agINs. Moreover, the circuit receives descending supraspinal signals.

**Figure 6.**
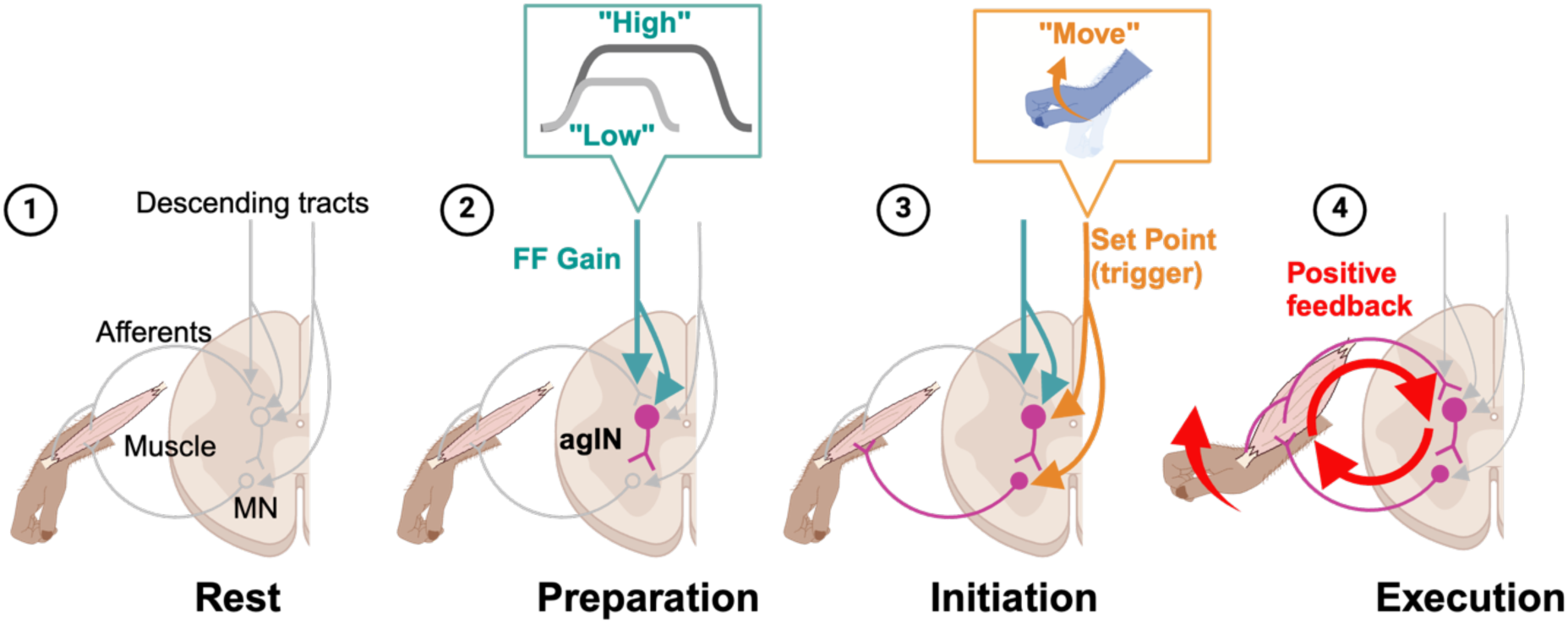
A positive feedback loop mediated by agINs and their mode of action for generating muscle activity during voluntary movement. Simplified schematics of the positive feedback loop mediated by agINs in different modes of action for generating muscle activity. ① *Rest*. No loop activity is observed during this stage. ② *Preparation*. The excitability of agINs starts to increase with input from the descending motor command (green arrows), and some agINs may exhibit spiking activity (see Extended Data Fig.6). However, the excitability of the target motoneurons and muscles remains below the threshold of activation, and thus, no movement initiated. The excitability of agINs at this stage reflects the duration and magnitude of the prospective EMG of their target muscles encoded by the descending tract command through increased feedback gain (top, an inset). ③ *Initiation*. The descending motor command sends a trigger signal to either agINs or their target motor neurons that is strong enough to bring their excitability to a supra-threshold, and as a consequence, EMG activity is generated. ④ *Execution*. The EMG activity generated in in the initiation state produced a joint torque and force feedback signal (red arrow). Consequently, they recurrently activate agINs and their target muscles (red circular arrows) until the duration and magnitude are reached to the predetermined level during the preparation state. Note that no descending inputs are required in the operation of the *execution* state.

During the motor preparation period (Fig. 6-②), the magnitude and duration of prospective motor action were set within the reflex loop at the agIN level (blue line, descending tract input), which functions as an FFG (blue lines, Fig. 4a). It is widely recognized that neurons in the primate primary and premotor cortices exhibit premotor activity, which is correlated with the performance of the upcoming movement^40–42^. These cortical pre-movement activities, encoding future spinal reflex’s states^43^, may be represented in the agINs through corticospinal input^44,45^, given the significant pre-movement activity observed in some agINs (7/15; Extended Data Fig. 6). In the spinal cord, the corticospinal neuronal activity can presynaptically and postsynaptically change the FFG to the agINs^4^. We recently reported that proprioceptive afferent input is facilitated (i.e., FFG increase) before and during agonistic voluntary movement^46^ owing to decreased presynaptic inhibition.

The transient ‘go’ signal from the corticospinal input^47^ triggers the looping activity (Fig. 6-③, orange lines) by activating either agINs^4^ or the motoneuron that is innervating target muscle^34,48,49^, which serves as a set-point input (Fig. 4a). The duration and magnitude of prospective motor action are already set within the loop (green lines); looping activity that is triggered (Fig. 6-④) can automatically generate, sustain, and terminate muscle activity without continuous regulation from the corticospinal tract.

These results prompted us to update our current understanding of the roles of the sensorimotor cortex and spinal reflex circuit in controlling muscle activity during voluntary movement. For decades, the neural control of volitional movement has been understood as a predominance of cortical control, especially via the corticospinal tract, based on abundant experimental data in monkeys and humans^12,50–52^. Here, we suggest that the spinal reflex loop operates as a self-sustaining unit that generates muscle activity with minimal involvement by descending commands (except setting the feedback gain and triggering looping activity). How can motor control mediated by the cortical and spinal circuits be orchestrated to perform complex motor actions in our daily lives? We suggest that, though both may have discrete roles, they can be unified by their hierarchical interactions. We assumed that, if the reflex gain is pre-programmed, spinal circuits can independently generate a relatively simple muscle-activity pattern (gross magnitude and duration). Supporting this assumption, our recent report indicated that spinal premotor neurons exhibit target muscle-like activity, whereas corticomotoneuronal cells diverge from this pattern in monkeys performing grasping movements ^33^.

However, in real-world situations, animals must interact with uncertain environments through intermittent, precise refinements of muscle activity. Such interactions are unlikely to be achieved by the proposed spinal reflex circuit due to the pre-programmed magnitude and duration of future muscle activities. We assume three major roles for descending systems. First, the sensorimotor cortex can function as a predictor^53^ of the action of the reflex loop. Herein, the crucial role of the sensorimotor cortex is to set the loop parameter (i.e., FFG) for upcoming movements and not to generate muscle activity *per se.* Using an updated prediction for the optimal parameter based on the prediction error from previous experience, the loop parameter was updated. Second, and most obviously, the spinal reflex loop needed to be triggered upon movement initiation (corresponding to the ‘set-point’ input in Fig.4A), and the corticospinal projection^12,49,54^ should have a predominant role as the trigger. Third, the cortical mechanism could directly, and not via by the spinal reflex system, modulate the activity of motoneurons and muscle activity^32^. This should be the most effective method for the online correction of muscle activity in skilled motor output.

Based on their gene expression profiles, spinal interneurons have recently been categorized into distinct classes,^55,56^ wherein the subtypes Dl3 and V2a exhibited characteristics similar to those of AgINs. Notably, both Dl3 and V2a INs are excitatory and project to the ipsilateral motoneuron pools. The Dl3 subtype plays a crucial role in forelimb grasping^57^ and hindlimb locomotion^58^. Genetic silencing of the dI3 IN output in mice caused loss of grip-strength modulation in response to increasing loads, thus mirroring the role of agINs in maintaining muscle force during hand movements in monkeys (Figs. 4 and 5). In contrast, a subset of Chx10-expressing V2a INs stabilizes and provides ipsilateral hindlimb locomotor activity^59^, and their ablation disrupts forelimb-reaching functionality^60^. Vieillard et al^61^ recently revealed that a subset of V2a INs receives sensory input from both the muscle spindle and Golgi tendon organs. This finding was consistent with the characteristic features of agINs, as illustrated in Fig. 1 and Extended Data Fig. 2. Though these spinal IN classifications have predominantly focused on rodents and other lower-level animals, bridging the gap between the genetic neuronal characteristics in mice and their roles in regulating voluntary movements in primates has immense potential for advancing our understanding of forelimb motor control.

## METHODS

### Animals

All experiments were approved by the Institutional Animal Care and Use Committee of the National Institute for Physiological Sciences (NIPS), Okazaki, Japan. We obtained data from three *Macaca fuscata* monkeys [KO and OK (female), and IS (male)]. During the training and recording sessions, each monkey sat upright in a primate chair with its right arm restrained and elbow bent at 90°. The hand of the monkey was held in a cast with the fingers extended and the wrist in the mid-supination/pronation position. The cast holding the monkey’s hand was attached to a servo-motor-driven manipulator that measured the flexion–extension torque about the wrist. The left arm was loosely restrained in a chair^19,20,62^.

### Behavioural paradigm

The monkeys performed a wrist flexion–extension task with an instructed delay period^19,20,63,64^ (Extended Data Fig. 1). The wrist flexion–extension torque applied to the spring-loaded manipulandum controlled the position of the cursor displayed on a computer monitor in front of the monkey. As the monkeys performed the task with their right hand, wrist flexion led to a leftward displacement of the cursor. Trials began with the monkey holding the cursor in a central target window corresponding to zero torque for 0.8 s. Next, the flexion and extension targets (empty rectangles) were shown to the left and right of the central target. One target flashed briefly for 0.2–0.5 s (Cue), indicating the correct movement to be performed at the end of the instructed delay period, signalled by the disappearance of the central target (GO). Trials were accepted only if no wrist movements occurred during the delay period (1 s). Following the GO signal, the monkey quickly moved (Active Movement) the cursor to the desired target (<1.5 s, including reaction time) and held the cursor in the target window for 1.5 s (Active Hold). The movements were performed against an elastic load applied using a servomotor (5 N*m). At the end of the Active Hold period, the peripheral target disappeared and the central target reappeared (Release GO). The monkey then relaxed its forearm muscles, allowing the servo spring to passively return the wrist (Passive Movement) to the zero-torque position (Rest). After keeping the cursor within the central target for 0.8 s, the monkey was rewarded with apple sauce (Reward) for successful trials.

### Surgical procedures

Following behavioural training, surgeries were performed aseptically after placing the animals under 1.5–3.0% sevoflurane anaesthesia (2:1 O_2_:N_2_O). Head-stabilization lugs were cemented to the skull with dental acrylic and screw-anchored to the bone. A recording chamber made by stainless-steel or polyetherimide resin (Ultem, Saudi Basic Industries Corporation) was implanted over a hemilaminectomy in the lower cervical vertebrae (C4–C7). Pairs of stainless steel wires (AS631, Cooner Wire) were implanted subcutaneously in 10–12 muscles [extensor carpi ulnaris (ECU), extensor carpi radialis (ECR), extensor digitorum communis (EDC), extensor digitorum-2,3 (ED23), extensor digitorum-4,5 (ED45), flexor carpi radialis (FCR), flexor carpi ulnaris (FCU), flexor digitorum superficialis (FDS), palmaris longus (PL), pronator teres (PT), abductor pollicis longus (APL), and brachioradialis (BRD)] that were active in one or both directions (i.e., flexion or extension). Each muscle was identified based on its anatomical location and the characteristic movements elicited by low-intensity intramuscular stimulation. Nerve-cuff electrodes ^65^ were implanted in the muscle branch (deep radial nerve) and cutaneous branch (superficial radial nerve) of the radial nerve, as well as in the median nerve^20^.

### General recording procedures

We penetrated the spinal cord dorsally through an implanted recording chamber with glass-insulated Tungsten or Elgiloy microelectrodes (impedance 0.8–1.4 MΩ; Fig. 1A)^19^. One to three penetrations were performed on each day of recording. When we encountered neurons exhibiting responses to DR stimulation (within the observation window of 10 ms), we measured the segmental latency using a landmark of the intraspinal volleys^20^. Neurons were identified as candidate first-order INs if the response occurred within 5 ms of the stimulus (Fig. 1B). We then asked the monkeys to perform wrist flexion–extension tasks and recorded the firing-rate modulation as a function of the task epoch for both the flexion and extension trials. For task performance, we examined the output of motoneurons projecting to the wrist flexor and extensor muscles (Fig. 1C) by STA^21,22^ (Fig. 1D). We identified the recorded spinal neurons as INs by confirming whether they (1) received the putative monosynaptic *input* from DR afferents and (2) sent an *output* to the motoneurons of the wrist and finger muscles by STA. Confirmation of the input–output profile of the recorded neurons was partially performed online but primarily completed post-recording.

### Identification of the reflex INs

#### Input to the spinal neurons

At the beginning of each electrode penetration, intraspinal volleys in the cord surface potentials were monitored (Fig. 1A), and the threshold currents that evoked an incoming volley from the DR were measured on most recording days. Subsequently, spinal IN single-unit responses were examined by stimulating DR with biphasic constant-current pulses (100 µs/phase) at a constant frequency of 1–2 Hz. The stimulus current was set at 1–1.2 times the threshold. Each action potential was isolated based on its waveform, and peristimulus time histograms (PSTH) were obtained for each IN. Segmental response latency was calculated from the first peak^66^ of the incoming volleys extracted from the cord surface potentials (average of all volleys; Fig. 1B, top) to the onset of the PSTH peak. We adopted a central latency of less than 1.5 ms as the criterion for putative monosynaptic linkage from the DR to neurons identified from each nerve^19,67,68^.

#### Output from the spinal neurons

The STA of the rectified EMG was computed to identify the output (post-spike effects, PSEs) of spinal INs on the recorded EMGs^22,69^. The STAs were compiled by averaging segments of rectified EMG activity from the 50 ms preceding each trigger to 50 ms after the trigger. Spikes (n>2000) were accepted as triggers only if the root mean square (RMS) value of the EMG from 30 ms before to 50 ms after the spike was greater than 1.25 times the RMS noise level in the EMG channel. STA was smoothed using a flat five-point finite impulse-response filter. The baseline trend was subtracted using the incremented-shifted averages method^70^, and significant STA effects were identified using multiple-fragment statistical analysis^71^ (*p*<0.00417 for 12 muscles, Bonferroni’s correction). The test window was set to 12 and 3–15 ms after the spinal neuron spike. Potential crosstalk between simultaneously recorded EMGs was evaluated by combining a cross-correlation method^35^ and a third EMG differentiation^72^, then STA effects potentially resulting from crosstalk between EMG recordings were eliminated from the present dataset. To distinguish the PSEs from the synchrony effects^73^, we measured the onset latency and peak width at half-maximum (PWHM); effects with onset latency >3.5 ms and PWHM <7 ms were identified as PSEs^22,69^. Neurons that showed a large ‘motor unit’ signature in the STA of the non-rectified EMG with only 50 spikes^74^ were identified as putative motoneurons and excluded from the data set.

Neurons confirmed with both monosynaptic input from DR nerve and presumed direct projection to the motoneurons were identified as the INs mediating spinal reflex (Reflex INs in Extended Data Fig. 2A). These neurons were classified according to their output patterns: the target muscles and effect of projection (Extended Data Fig. 2B). When the only target muscles of INs were the wrist extensors, they were classified as autogenic. When those were only wrist flexors, they were classified as heterogeneous. When those were both extensors and flexors, they were classified as both. In contrast, INs with positive PSE to the target muscles were classified as excitatory, whereas INs with negative PSE were classified as inhibitory.

### Decoding agIN activity by EMG signals

First, we preconditioned the EMG and agIN firing rates for the decoding analysis. For the EMG signals, we performed temporal filtering of the EMG signals using a second-order Butterworth high-pass filter (5 Hz). The EMG signals were rectified and computed in 1-ms bins. We calculated the smoothed curves for the signals using a mobile window process with 11 bins. For agINs, we first calculated the instantaneous firing rate by convolving the inversion of the interspike interval with an exponential decay function with a time constant of 50 ms. We computed the firing rate in 1-ms bins, corresponding to the resampling frequency of the EMG. We then applied a Bayesian Sparse Linear Regression (SLiR) algorithm that introduced sparse conditions for only the channel dimension, and not for the temporal dimension of the model. Using multidimensional linear regression, the instantaneous firing rate of spinal interneurons was modelled as a weighted linear combination of pre-processed EMG signals as follows:

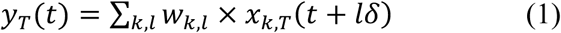

Where; y_T_(t) is a vector of activity of a spinal interneuron at time index t in a trial T. x_k,T_(t+lδ) is an input vector of an EMG signal k at time index t, and time-lag lδ (δ = 1 ms) in a trial T. w_k,l_ is a vector of weights on a peripheral afferent or a cortical electrode k at time-lag lδ, and b is a vector of bias terms to y_T_. As we examined how EMG signals influenced the spinal IN activity, time-lag lδ (Equation 1) was set to negative values. We used EMG signals from −50 ms to time −1 to decode the spinal IN activity at time 0. This shorter lag eliminates the potential involvement of long-loop reflexes^32^.

Subsequently, using a training dataset, we built a model to predict the firing rate of spinal IN and tested it using a test dataset. Twenty trials were randomly selected as the training dataset, and one other trial was selected as the test dataset. To assess the model, we calculated the correlation coefficient between the observed and reconstructed spinal IN activities in the test dataset. We performed a 10-fold cross-validation for the analysis of each session and used the average values for the analysis. In the control analyses of model reconstruction, we created surrogate training datasets wherein we randomized the temporal profile of the EMG signals to generate and subsequently test a model.

### Reconstruction of EMG signals from the PSE of spinal INs

To reconstruct the EMG from the PSE, we first extracted a 15-ms STA time-series from the spike onset as a template (shaded area in Fig. 3B-a). Next, we calculated the peri-event time histogram (PETH) of each agIN from −1 to 3 s from movement onset with a 5-ms bin and calculated the convolution of the PETH and template (Fig. 3B-b) to obtain the reconstructed EMG, which was then filtered using a 10–200 Hz bandpass filter (Fig. 3B-d); the onset and offset of muscle activity were defined with a successive overshooting two-fold standard deviation of the baseline. Finally, we calculated the correlation coefficient (Fig. 3C) between the real EMG (Fig. 3B-c) and the reconstructed EMG (Fig. 3B-d).

### Positive feedback model

#### Diagram

We constructed a modified version of a nonlinear model of the cat hindlimb segmental reflex system^39^. In this model (Fig. 4A), the EMG signal at α-motoneuron is generated by force- and displacement feedback in the closed-loop configurations. An independent drive to the α-motoneuron from outside the closed-loop, emulating the descending motor command (‘set-point’), was allocated. To initiate the closed-loop activity, we applied a triangular pulse (200 ms) to the set point (Fig. 4B, top) and examined the time-dependent changes in the EMG signal as an outcome of the closed-loop activity (black line in Fig. 4B bottom). We then compared these with the EMGs reconstructed using each agIN (grey in Fig. 4B, bottom). In this simulation, we have three variables – the FFG (Ib gain, blue in Fig. 3A), the displacement feedback gain (Ia gain, orange in Fig. 3A), and the amplitude of set-point (red in Fig. 3A). We explored the optimal set of the three variables that provided the best match between the simulated and reconstructed EMG signals. In this procedure, the beta gain was fixed to illustrate the effects of changing a given variable.

The model consists of the muscle model using a modified Hill model^75^, displacement feedback via muscle spindle Ia afferents, force feedback via tendon organ Ib afferents, positive feedback via the β-fusimotor pathway, and physical environments, as described previously^39^. Except for the gain parameters, we used the same parameters as the original model^39^ (Extended Data Table 1). The physical environment (the hand of the monkey) was constructed as a one-link model with a damper.

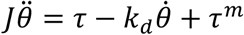

The wrist as the axis ^76^. Where *θ* is the angle of wrist, *τ* is the joint torque generated by the muscle, *τ^m^* is the physical effect of the manipulandum. Moment of inertia *J*=6.38×10^-3^ kgm^2^ ^77^, viscosity of wrist *k_d_*=0.06 Nm s/rad were respectively set as fixed values based on the –

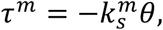

and the spring constant 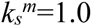 Nm/rad was determined from its mechanical property.

### Exploration of the optimal network parameter for simulating ag-IN generated EMGs

To find the FFG gain, displacement-feedback (DFG) gain, and set-point amplitude (SP) that correspond to the reconstructed (Fig. 3) or actual EMG data (Extended Data Fig. 5), we compute the RMSE (Root Mean Squared Error) between the simulated EMG signal (α-motoneuron output) and the reconstructed or actual EMGs, and search for the FFG and SP that minimize the RMSE. Here, the DFG for optimization was set to 0.5×FFG (Fig. 4G, J and Extended Data Fig. 5G, J support that the optimal result is not sensitive to the DFG value and that this assumption is reasonable). To determine the optimal parameters, we used the interior-point method with non-negative constraints. We used MATLAB ‘fmincon’ function for the implementation.

#### Examining the effect of FFG, DFG, and SP

To reveal the influence of the FFG, DFG, and SP on the simulated EMGs, we gradually changed each from their optimal values and performed a simulation for each parameter (Fig.3C-E and Extended Data Fig.5 C-E).

#### Characterization of the simulated EMG signals

To facilitate a comparison of simulated EMG across different FFG, DFG, and SP, which were systematically varied around their optimal values for the 22 agIN-muscle pairs, we characterized the simulated EMGs using two key metrics: normalized amplitude and sustaining duration of simulated EMGs (Fig. 3F–M and Extended Data Fig. 5F–M). The normalized sustained duration was calculated as follows:

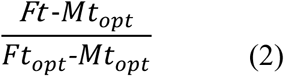

where Ft is the time at which the simulated signal falls below half of the maximal amplitude, Mt is the time of the peak amplitude, and opt is the optimal value. The normalized amplitude was calculated as follows:

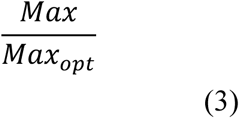

where Max is the peak amplitude and Maxopt is the peak amplitude in the simulated EMGs obtained using the optimal parameter.

### Classification and characterization of successful and error trials

Each trial was considered successful if the monkey completed all the task epochs. Any trial lacking at least one of the task epochs was defined as an erroneous trial. The error trials were classified into three types: (1) *No movement*, trials without any detectable changes in torque after the go signal was delivered. These trials were aborted after the grace period (0.2–0.5 s after the go signal); (2) *Wrong direction*, trials in which movement after the go signal was opposite from the instructed direction; (3) *Short-hold*, trials in which the monkey moved in the correct direction, but did not hold the instructed wrist torque long enough. The trial was aborted when the torque exceeded or decreased the determined range without returning to the correct range within 0.1 s.

We characterized the EMG signals of the target muscles during successful and erroneous trials by measuring the amplitude and duration of each wrist extension trial. To this end, the EMG of the target muscle was bandpass filtered (50–200 Hz) and rectified. The rectified EMG was again filtered using a 5-Hz low-pass filter to emboss the offset of EMG. The onset of EMG was defined as the first point that exceeded the time of the two-fold SD from the average value at the remaining epoch. The offset of the EMG was defined as the local peak where the value dropped the most sharply at the next local peak. The mean amplitude and duration of the EMG for each trial were measured using the EMG onset and offset time points.

### Electrophysiological evaluation of afferent feedback gain in each agIN and their relationship with the EMG amplitude and sustained duration of future wrist movement

We stimulated the DR with biphasic constant-current pulses (100 µs/phase) at a constant frequency of 1–2 Hz during the task and measured the evoked response of each agINs during the rest, cue, and delay period in the wrist extension trials (both successful and erroneous trials). For each agIN, the mean evoked response was computed by pooling all the stimulation pulses applied during the task epochs to maximize the signal-to-noise ratio. We then used the onset and offset points of the mean evoked response to compute the peak area evoked in each task period (i.e., using the stimulation pulses applied during a specific period). The peak area represents the firing probability of each agIN for DR stimulation with comparable stimulus intensity. We define this peak area as the input-output gain of each agIN, as described previously^20^.

We performed a trial-based analysis to estimate the trial-by-trial alternation of the relationship between the afferent input gain of each agIN and the amplitude and duration of the EMG of the target muscles. For this analysis, we examined whether electrical stimulation elicited a response of agIN activity (spike) within 5 ms of stimuli application during the rest, cue, or delay periods in the wrist-extension task (both successful and erroneous trials). Two or three stimuli were typically applied during task epochs. If one of these stimuli elicited the response of agINs, then we defined the trial as the “effective,” otherwise as the non-effective” trial. Thus, all trials performed during the recording from each agIN, in which at least one stimulus was applied during these task epochs, could fall into two categories.

Next, we measured the maximal amplitude and duration of the EMG of each agIN’s target muscles during ‘effective’ and ‘non-effective’ trials. All EMG signals were bandpass-filtered (50–200 Hz), rectified, and low-pass-filtered. The onset of EMG bursts that preceded the movement onset was defined as the starting point, with signals exceeding two standard deviations above the mean EMG amplitude during the rest epoch for at least 50 ms. The offset of EMG bursts that preceded movement offset (defined as the timing of movement onset +1.8 s) was defined as the final point, with signals exceeding one standard deviation above the mean EMG amplitude during the rest epoch for at least 200 ms. The EMG duration was defined as the difference between the timing of EMG onset and offset. The mean EMG amplitude was defined as the mean rectified EMG amplitude between the onset and offset. Subsequently, for each recording day, we sorted the corresponding trials by the mean amplitude (from smaller and larger) or the duration (from shorter and longer) of the EMGs burst and numbered them. For normalization to compile the data on different days, each number was then divided by the number of trials per day. Finally, we compared the distribution of these ‘relative amplitude’ and ‘relative duration’ of the EMG burst of the target muscle between ‘effective spikes’ and ‘non-effective’ trials by making histograms (bin size=0.1).

### Statistical analysis

All parametric and nonparametric tests used, as well as any multiple-comparison tests and corrections, are indicated in the figure legend or main text, as is the *n*-value for each analysis. All statistical comparisons were two-tailed, when relevant. *P*<0.05 was considered significant, and * indicates *p*<0.05 and ** indicates *p*<0.01; all significant *p*-values are indicated in the figure legends. Statistical analyses were performed using MATLAB or Prism version 9.2.0 (GraphPad Software, San Diego, CA, USA).

## Data availability

All data are available in the main text, the Extended Data materials.

## Acknowledgments

We would like to express our sincere gratitude to Drs. Ole Kiehn and Gerald E. Loeb for their invaluable comments on the earlier version of the manuscript. We would also like to thank Nobuaki Takahashi and Kaoru Isa for their technical assistance. This work was partially supported by Grants-in-Aid JP18020030, JP 18047027, and JP 26120003 from the Ministry of Education, Culture, Sports, Science, and Technology of Japan (MEXT; to K.S.) and the Precursory Research for Embryonic Science and Technology program (JPMJPR09G8) from the Japan Science and Technology Agency (JST; to K.S.). This work is also supported in part by the NSF CRCNS Japan-US 2113096 to K.S. (Subaward PI) and Francisco Valero-Cuevas (PI).

## Author Contributions

K.S. designed the study. J. K., T. T., and K. S. performed the experiments. J.K., S.T., T.U analysed the data. T.F. and K.S. designed and conducted the simulations. K.S. wrote the manuscript. All authors have approved the manuscript.

## Author Information

The authors declare that they have no competing financial interests. Correspondence and requests for materials should be addressed to K.S.

---

**Extended Data Figure 1.**
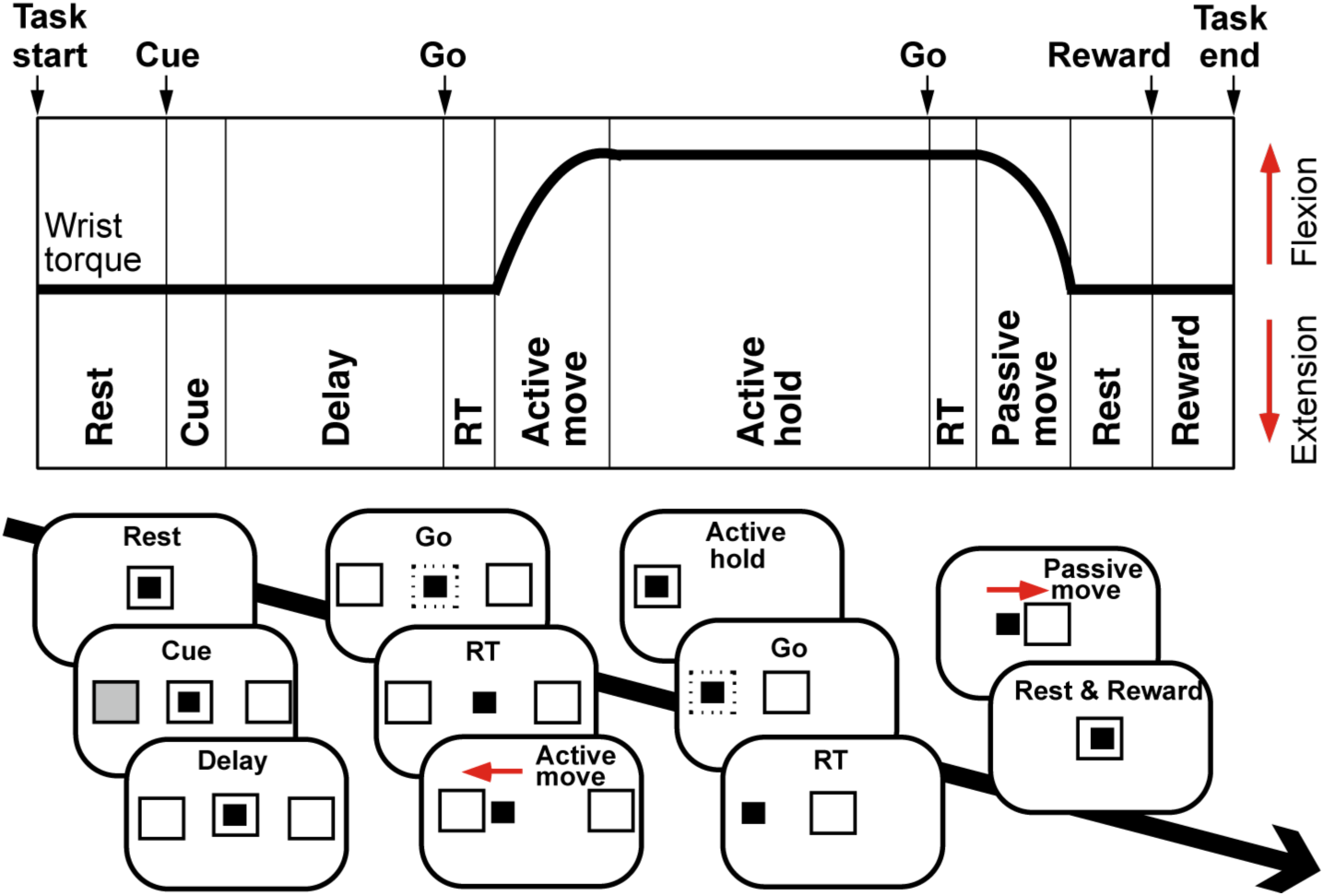
Whist flexion–extension task with an instructed delay period. Top: Schematic illustration of wrist torque measured during a single flexion trial. Bottom: Display instructions and presentation of monkey wrist movements. The black-filled square represents a moving cursor, the solid empty squares represent central and peripheral targets, and the grey-filled square represents the peripheral cue. The red arrows indicate the direction of movement. RT: reaction time. During the flexion trials, the active movement and hold epochs required a wrist flexion movement, and the passive movement involved wrist extension (and vice versa for extension trials). See Methods for further details.

**Extended Data Figure 2.**
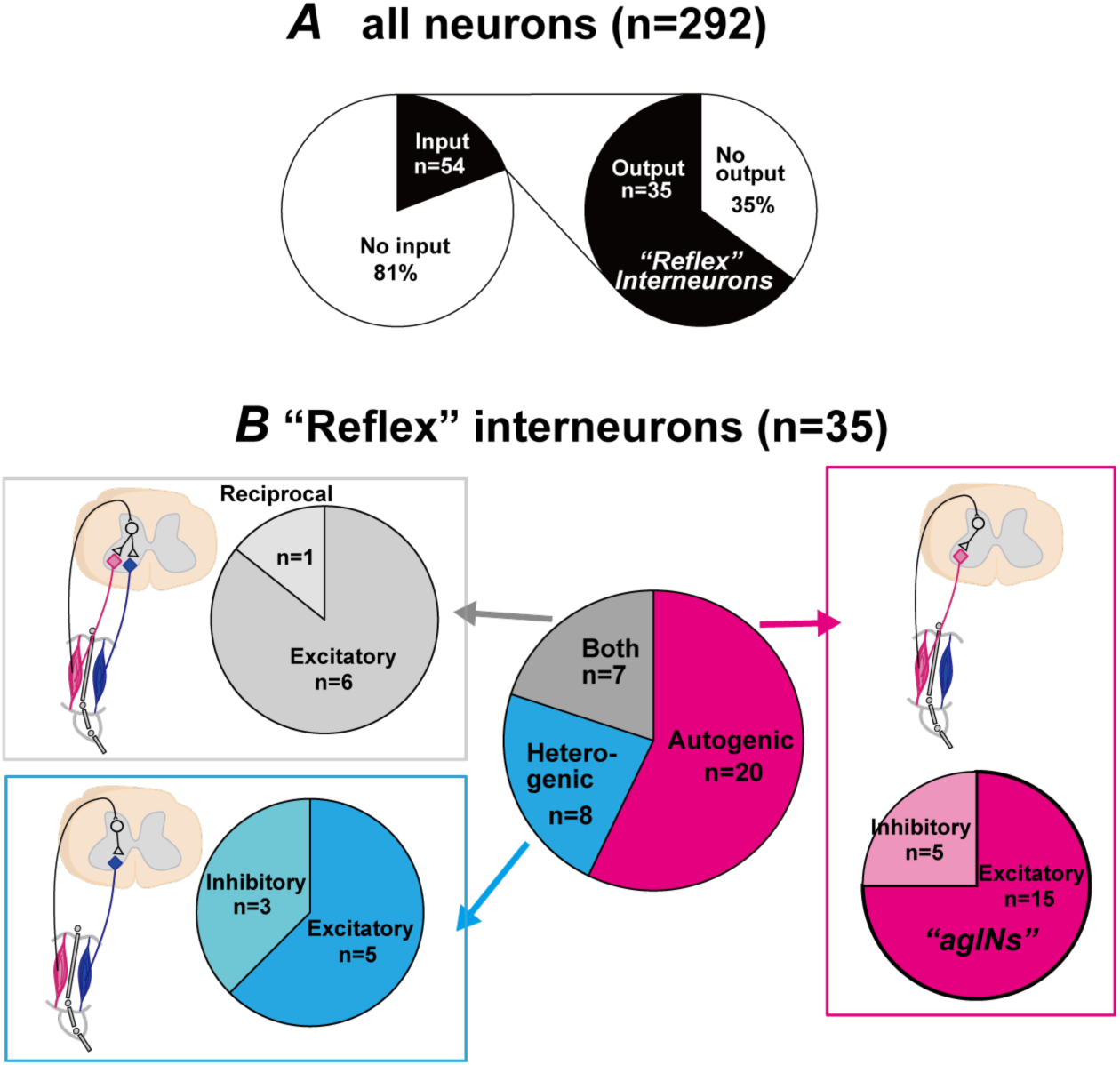
Population of spinal interneurons recorded in this study. **A**: A total, 292 neurons were recorded from three monkeys, and 54 (13.8%) showed input from the DR nerve within a central latency of 1ms. Among these, 35 exhibited a post-synaptic effect after spike-triggered averaging. **B**: Classification of reflex-related INs by their input-output pattern. Autogenic: The output is directed to the extensor muscles. Heterogenic: Output of the flexor muscles. Both: Output to flexor and extensor muscles. Excitatory: INs with post-spike facilitation (e.g., excitatory INs). Inhibitory: INs with post-spike suppression (e.g., inhibitory INs). The 15 excitatory INs exhibiting autogenic input-output patterns (agINs) are the main focus of this study.

**Extended Data Figure 3.**
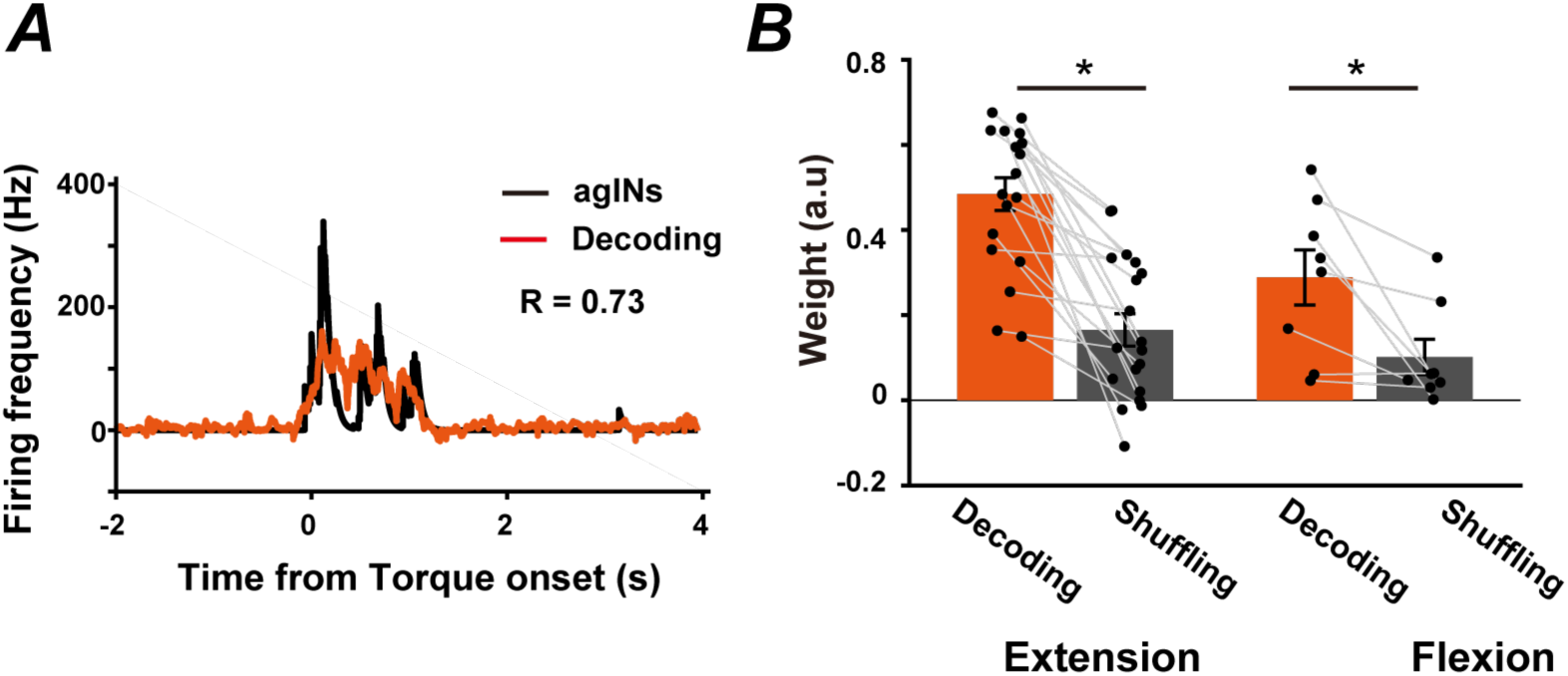
Decoding the agIN’s firing profiles by the EMG. **A**: An example of the reconstruction of an agIN using the EMGs of the forelimb muscles of Monkey O. Black, actual activity; red, reconstructed activity. R: Correlation coefficient between observed and reconstructed activities. R=0.73. **B**: Mean (±SEM) decoding accuracy pooled across INs. Correlation coefficients between the actual and reconstructed traces are presented (\**P* < 0.05, n=19 for extension, n=8 for flexion).

**Extended Data Figure 4.**
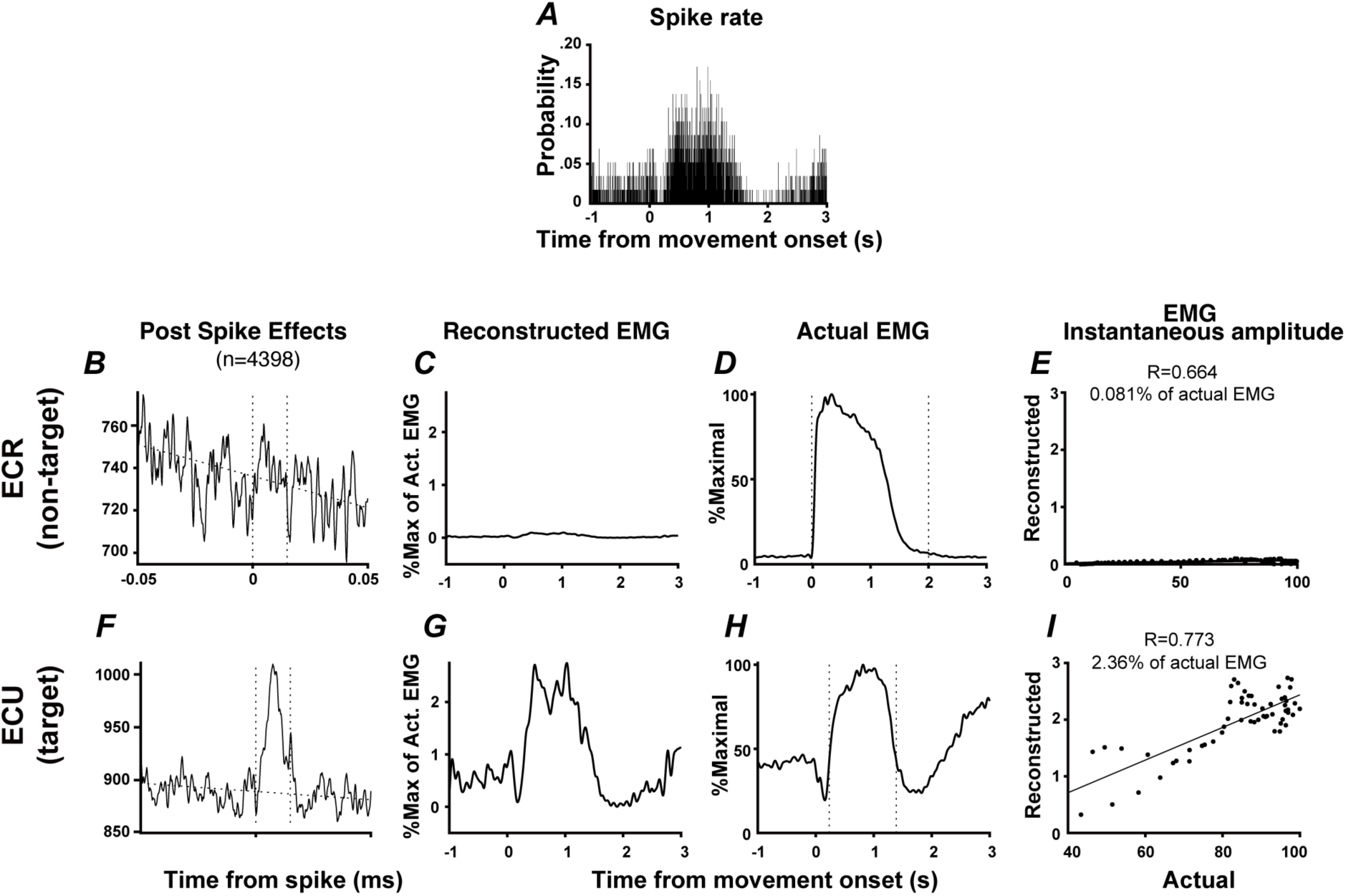
Comparison of original- and reconstructed EMGs of target and non-target muscles of single agINs. The second example of the actual- and the reconstructed EMG by convolution of the snippet of post-spike effects on the target- and non-target muscle to the spike timing of single agINs. (**A**) P.S.T.H of single agIN aligned to the movement onset of extension trials (n=15). MO, movement onset. (**B-E**) Post-spike effect of agINs shown in A (B), the reconstructed EMG (C), actual EMG (D) and the EMG instantaneous amplitude and linear regression line (E) for the non-target muscle. R, correlation coefficient. (**F-I**) Same format but showing results for the target muscle of agINs.

**Extended Data Figure 5.**
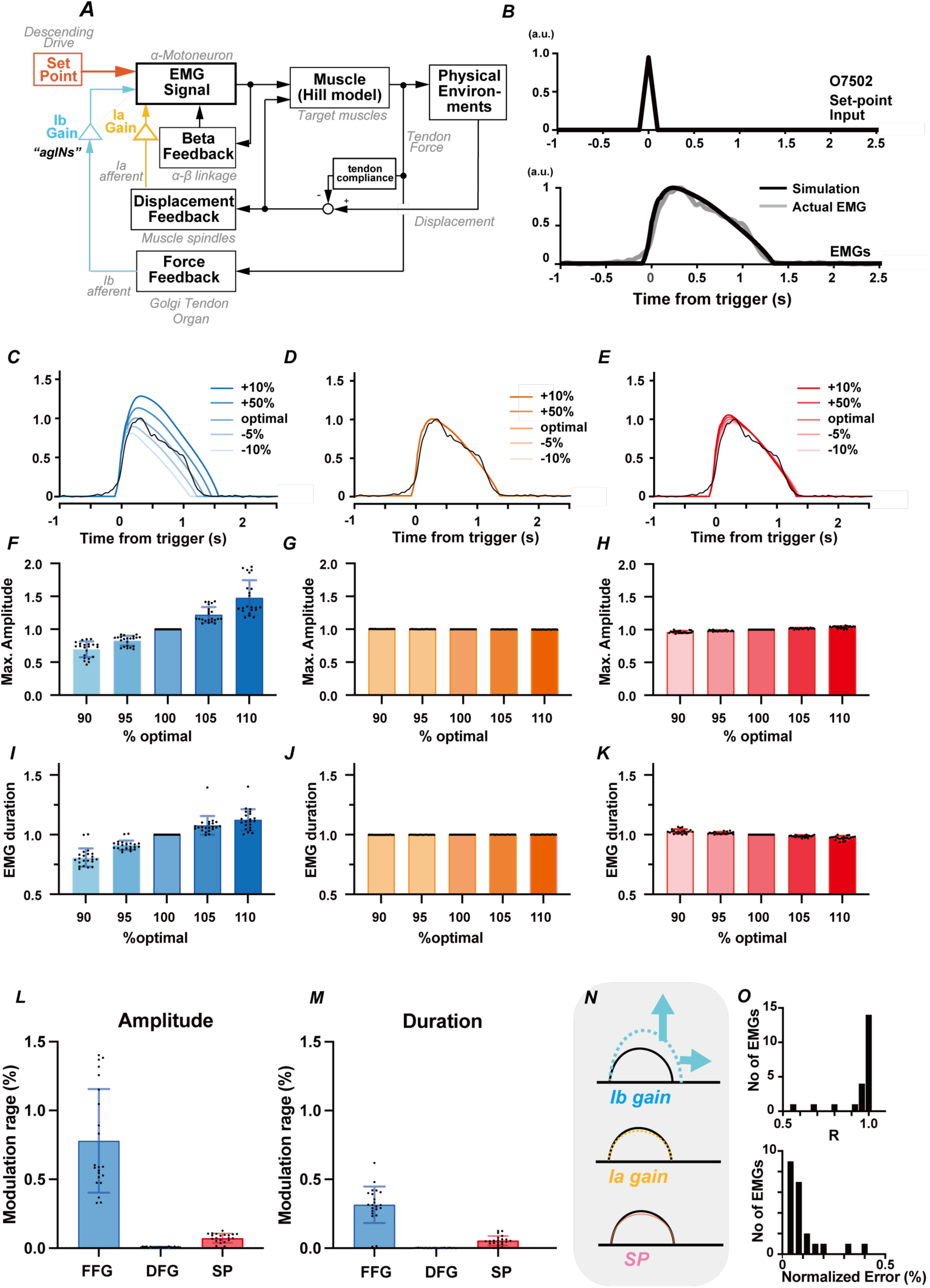
Simulation for actual EMGs. Result of simulation for the actual EMG. See the text and legends for Fig. 4.

**Extended Data Figure 6.**
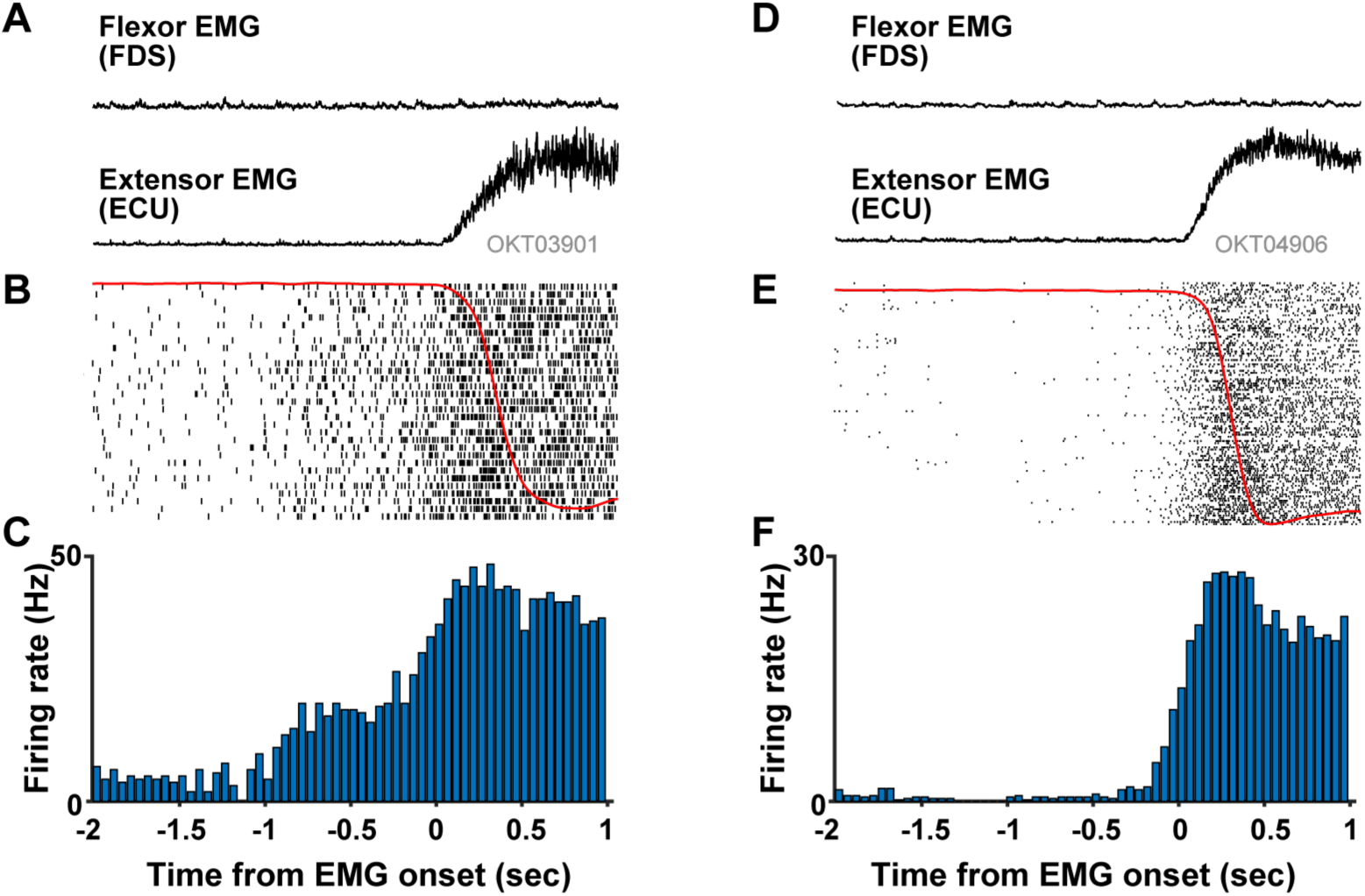
Pre-EMG activity in agIN. Examples of agIN spiking activity shown with simultaneously recorded EMG signals in wrist extension trials. The agIN activity and EMGs signals were aligned with the onset of the extensor EMG signal. (**A,D**) EMG of the flexor (FDS) and extensor (ECU) muscles. (**B,E**) Raster plots showing the activities of single agINs. The red line overlaying the plot represents the average wrist torque. (**C,F**) Peri-stimulus time histograms quantifying the rate of modulation of agINs starting before the onset of EMG. Two examples of agIN with (**A-C**, n=32 trials) or without (**E-F**, n=114 trials) pre-EMG activity are shown.

**Extended Data Table 1.**
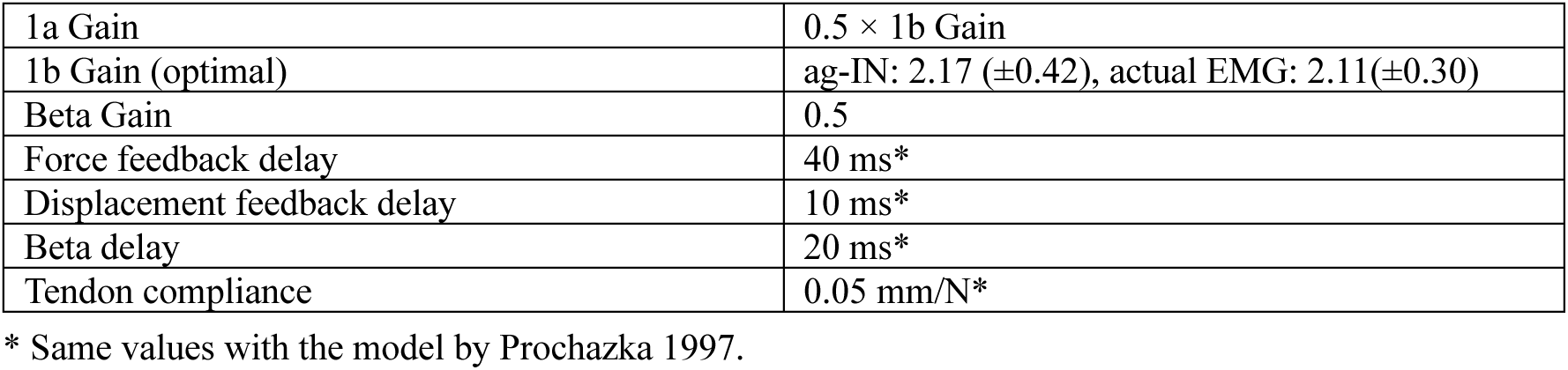

